# Multi-modal Efficacy of a Chimeric Vesiculovirus Expressing the Morreton Glycoprotein in Sarcoma

**DOI:** 10.1101/2022.09.18.508434

**Authors:** Chelsae Dumbauld, Oumar Barro, Natalie M. Elliott, Yumei Zhou, Musa Gabere, Elizabeth Raupach, Alexander T. Baker, Michael T. Barrett, Kenneth H. Buetow, Bertram Jacobs, Mahesh Seetharam, Mitesh J. Borad, Bolni Marius Nagalo

## Abstract

Vesiculoviruses are attractive oncolytic virus (OV) platforms due to their rapid replication, genome with appreciable transgene capacity, broad tropism, limited pre-existing immunity, and type I interferon response gradient between malignant and normal cells. We developed a synthetic chimeric virus (VMG) expressing the glycoprotein (G) from Morreton virus (MorV) with remaining genes from vesicular stomatitis virus (VSV). VMG exhibited *in vitro* efficacy in cell proliferation and cell death assays across a broad range of sarcoma subtypes and across multiple species. Notably, all cell lines tested showed ability of VMG to yield productive infections with rapid replication kinetics. Pilot safety evaluations of VMG in immunocompetent, non-tumor bearing mice showed absence of toxicity with intranasal doses as high as 1e^10^ TCID_50_. VMG resulted in tumor reduction *in vivo* in an immunodeficient subcutaneous Ewing sarcoma model at doses as low as 2e^5^ TCID_50_. In the immune competent murine syngeneic fibrosarcoma model, while no tumor inhibition was achieved with VMG, there was a robust induction of CD8+ T cells suggestive of potential for combination approaches with immunomodulatory agents such as immune checkpoint inhibitors. The studies described herein establish the potential for VMG as a novel oncolytic virotherapy platform with multi-modal anti-tumor effects in sarcoma.

## Introduction

Sarcoma is a heterogenous disease, arising from non-epithelial tissues including bone, fat, cartilage, and muscle.^1^ Current treatments for advanced sarcomas comprise of chemotherapy, radiation, and surgical resection which are associated with both short- and long-term side effects.^2^ Owing to vast heterogeneity in responses, the 5-year survival rate for advanced stage soft tissue sarcoma or bone sarcoma is less than 20%, emphasizing the need for novel therapeutic strategies for treatment of refractory sarcoma.

Oncolytic virotherapy has recently emerged as a promising approach and offers an opportunity for synergistic combinations with other established cancer therapies such as immune checkpoint inhibitors.^1,3,4^ VSV is a particularly attractive oncolytic due to its broad tropism, low seroprevalence in the general population, ease of genome manipulation, and favorable carrying capacity for transgenes. However, limitations to VSV-based therapies include safety concerns based on this tropism, which has evidence of inducing neurotoxicity in murine and Rhesus macaque models.^6^ Current clinical candidates of VSV-based therapy include an IFNβ transgene to increase safety of the virus. Vesicular stomatitis virus (VSV)-derived vectors have advanced to phase II of clinical testing in cancer across a spectrum of hematological and solid tumor malignancies (NCT04291105).^5^ Another barrier for VSV therapy is humoral immunity, which generates neutralizing antibodies mainly targeted to the viral glycoprotein (G protein).^7^ These neutralizing antibodies can abrogate the effect of repeat virus administrations.

While there has been evaluation of several oncolytic virus platforms preclinically and in human trials against multiple epithelial cancers, there has been limited investigation of OVs in sarcomas. We therefore sought to evaluate oncolytic potential of novel chimeric vesiculoviruses in pre-clinical sarcoma models.

We and other groups have explored the use of attenuated or alternative vesiculoviruses such as Morreton virus (MorV) in the treatment of cancer.^8,9^ We have previously demonstrated that MorV exhibits a lack of neurotoxicity and hepatotoxicity in contrast to the concerns associated with VSV.^9^ In addition, we have shown that intratumoral injection of MorV elicited a cytotoxic T-cell response resulting in a prominent tumor regression at a 10-fold lower dose than VSV in a syngeneic cholangiocarcinoma model. In the current study, we examined the potential for use of a chimeric vesiculovirus in sarcoma, using the genetic backbone of VSV expressing the glycoprotein of MorV (VMG). With this approach, we hope to leverage the existing data derived from clinical studies on VSV-based vectors in combination with the pre-clinical safety profile of MorV with the ultimate goal to translate VMG into the clinic.^9^

## Materials and Methods

### Recombinant Virus Generation

Plasmid pVSV-XN2 was a gift from Richard Vile (Mayo Clinic Rochester, USA) encoding for full-length antisense VSV Indiana genome. Wild type Morreton virus was obtained from the University of Texas Medical Branch (UTMB). Full length MorV genome was confirmed by RNA-sequencing at the Iowa State University Veterinary Laboratory (ISU VDL). DNA sequence encoding for full-length MorV G polypeptide and the intergenic regions (IGRs) between MorV M and G genes (25 nucleotides) and MorV G and L genes (41 nucleotides) were synthetized by Genscript (Piscataway, NJ). Using laboratory cloning methods, VSV-G gene was removed from plasmid pVSV-XN2 and replaced by that of Morreton virus (MorV-G) G gene and its IGRs to produce pVSV-MorV-G. To rescue infectious VSV-MorV-G particles, BHK-21 cells were infected with vaccinia virus (Kerafast; Boston, MA) expressing the T7 RNA polymerase. Vaccinia infected BHK-21 cells were transfected with plasmids encoding for full-length pVSV-MorV-G and pVSV-L, pVSV-N, pVSV-M and pVSV-P genes as described by Lawson et al.^12^ Cell supernatant containing infectious VSV-MorV-G (VMG) was collected and filtered through 0.22 μm filter (MilliporeSigma; Burlington, MA) to remove residual vaccinia virus. VMG filtered supernatant was used to infect fresh BHK-21 cells until cytopathic effect (CPE) was apparent under light microscope. Complete genome sequence of VMG containing VSV N, M, P and L genes along with MorV G gene and IGRs were confirmed by ISU VDL using RNA-sequencing.

### Cell Lines

A673, SK-LMS-1, SK-UT-1B, Sarcoma 180, WEHI 164, 143B, and BHK-21 were obtained from American Type Culture Collection (Manassas, VA). OSCA-71 was purchased from Kerafast (Boston, MA). Cells were maintained in appropriate cell culture medium as directed by supplier with 10% heat-inactivated fetal bovine serum (FBS) (R&D Systems; Minneapolis, MN) and 1% Penicillin-Streptomycin (Thermo Fisher) in a 37°C, 5% CO_2_ incubator (HeraCell; Thermo Fisher).

### LDLR Flow Cytometry

Adherent cells were scraped from flask bottoms, and suspension cells were pipetted from flasks into round-bottom 96 well plates. Cells were washed with PBS, Fc receptors blocked, then stained with PE-conjugated mouse or human LDLR antibody. Cells were washed with 2% FBS in PBS and fixed in 2% PFA. Data was acquired on an LSR Fortessa (BD). Data was analyzed in FlowJo 10.8.1 FlowJo (RRID:SCR_008520) (BD). LDLR positivity was determined by a 5% isotype control gate on single cells.

### Viral Stock Production

Viral stocks were made by infection of T-225 flasks (Corning) of 80% confluent BHK-21 cells with MOI 0.001. When cytopathic effects were seen, media was collected from the flask and centrifuged to remove cell debris. Virus was purified using 10-40% sucrose density gradient ultracentrifugation followed by dialysis. Fifty percent tissue culture infectious dose (TCID_50_) values were determined by the Spearman-Kärber algorithm using serial viral dilutions in BHK-21 cells.

### Viral Growth Kinetics

2.5e5 cells were plated in each well of a 6-well plate in 2 mL DMEM 10% FBS 1% P/S. After overnight rest, cells were infected with VSV-MorV-G at MOI 10, 1, 0.1, and 0.01 or mock. After 1 hpi, cells were washed, and fresh media was added. At timepoints 2, 6, 24, 48, and 72 h, supernatant was taken and stored at −80°C. Viral titer was performed with serial dilutions of supernatant on BHK-21 cells. Fifty percent tissue culture infectious dose (TCID_50_) values were determined by the Spearman-Kärber algorithm.

### Human IFNβ ELISA

Quantikine Human IFNβ ELISA (R&D systems; Minneapolis, MN) was performed according to manufacturer’s recommendations on cell culture supernatants from sarcoma cells infected with MOI 10 and 1 collected at 24, 48, and 72 hpi which were stored at −80°C after collection.

### MTS Assay

Cell lines were plated at 1e4 cells/well in a 96-well plate and rested overnight in a 37°C/ 5% CO_2_ incubator. The next day, viral serial dilutions were added in three replicates per MOI. At 72 h, CellTiter96® AQ_ueous_ One Solution Cell Proliferation Assay (MTS) (Promega Corp, Madison, Wisconsin, USA) was added to each well as recommended by manufacturer and incubated for 1 h at 37°C. Absorbance at 490 nm was read using Cytation3 plate reader (BioTek; Winooski, VT). Viability was normalized to the maximum absorbance read per plate as representative of 100% and background absorbance was subtracted. A minimum of 3 experimental repeats were performed.

### Crystal Violet Assay

1e5 cells/well were plated in 12-well plates and rested overnight. The next day, virus was added at MOI 10, 1, and 0.1. At 72 hpi, cells were washed twice with cold PBS, fixed on ice in cold methanol, and then stained with 0.5% crystal violet in methanol. Plates were gently washed and dried overnight before taking phase images with EVOS FL Auto microscope at 10x magnification.

### IncuCyte Assay

A673 cells were plated at 1e4 cells/well in a clear 96-well plate (Falcon) and rested overnight. The next day, Annexin V Red dye was added to assay wells, followed by addition of 10-fold viral dilutions. Plates were sealed with Breathe-Easy membrane (Research Products International; Mt. Prospect, Illinois) and placed into IncuCyte S3 (Sartorius; Germany). Four phase and red fluorescence images per well were taken every 3 hours for a total time of 72 h. Images were analyzed in IncuCyte software for red object area. Data was exported to Excel and four images per well and three wells per MOI were used to graph data in GraphPad Prism v9 to generate sigmoidal, 4PL non-linear regressions. The calculated IC50 corresponded to the time at which 50% of maximal cell death had occurred (T_50_).

### Animal Studies

Animal work was approved by the Mayo Clinic Institute of Animal Care and Use Committee. All experiments conformed to regulatory standards. Mice were maintained in specific pathogen free environment.

### Toxicity Evaluation

VMG was intranasally administered at doses of 1e7, 1e8, 1e9, or 1e10 TCID_50_ in 10 μL to female Balb/c mice, with 6 mice per experimental group. Saline was administered intranasally in the control group. Mice were observed daily for adverse events including circling, paralysis, listing, seizure, or changes in behavior. Body weight was measured 3 times per week for a total of 49 days.

### Xenograft Model

Female NOD-SCID mice were subcutaneously inoculated with A673 cells into right flank. When average tumor volume reached >150 mm^3^ (Day 0), one dose of VSV-MorV-G was administered intratumorally at 2e5 or 2e6 TCID_50_ in 50 μL PBS or PBS only in the control group. Tumor dimensions were measured by digital caliper three times per week and tumor volume calculated. Mouse body weight and clinical observations were also recorded three times per week.

### Liver Function Tests

Liver function tests were performed by loading 150 μL of undiluted mouse serum into a Dri-chem 4000 chemistry analyzer (Heska; Loveland, CO) using the Heska liver function panel cartridges.

### Syngeneic Model

1e6 WEHI 164 cells in 100 μL 50% solution of Matrigel (Corning) were inoculated into the right flank of 24 male and female Balb/c mice with an average weight of 20 g. Tumor dimensions were taken by digital caliper measurements twice per week and tumor volume calculated. When tumor volume reached 80-120 mm^3^, mice received an intratumoral injection of 1e7 TCID_50_ VMG or 50 μL PBS (Day 0). Intratumoral injections were repeated at day 5 and day 10. The study was concluded at day 14.

### WEHI 164 Sample Collection

Mice were sacrificed at humane endpoint (tumor >10% body weight or ulcerated) or end of study. Blood was collected by cardiac puncture into clot tubes, placed on ice for 30 minutes. Serum was isolated by centrifugation and collection of supernatant, which was then frozen at −80°C. Tumors were excised, minced, then dissociated in 5 mL RPMI containing 1.5 mg/mL Collagenase II (MP Biomedicals; Irvine, CA) using GentleMACS C tubes on a GentleMACS dissociator (Miltenyi; Germany). Suspension was filtered through a 70 μM filter (Corning; Corning, NY) then washed twice with RPMI and cryopreserved in 10% dimethyl sulfoxide in FBS.

### WEHI 164 Immunophenotyping

Cryopreserved TILs were thawed in warmed RPMI plus benzonase and washed in RPMI. Cells were plated in a round-bottom 96-well plate at 1-2e6 cells/well. Cells were washed with PBS, blocked with mouse CD16/32 Fc block (BioLegend; San Diego, CA), then stained with Zombie UV dye (BioLegend), surface antibody cocktail, then permeabilized with eBioScience FoxP3 Fixation/Permeabilization kit (eBioScience, UK) according to manufacturer’s protocol. They were stained with intracellular antibody cocktail, then fixed with 2% paraformalin and kept on ice until acquisition on a BD Symphony A3 (Becton Dickinson BD; Franklin Lakes, NJ). Data was analyzed using FlowJo 10.8.1 (RRID:SCR_008520) (BD).

### IFNβ Sensitivity Assay

Cells were plated at 1e4 cells/well in a 96-well plate and rested overnight. Edge wells were loaded with media only to avoid edge effects. Serial two-fold dilutions of IFNβ were added across the plate and serial 10-fold dilutions of VMG were added down the plate. Plates were incubated for 72 h and then MTS was added to each well according to manufacturer’s protocol. After one hour, absorbances were read at 490 nm using Cytation3 plate reader (BioTek; Winooski, VT). Viability was normalized to the maximum absorbance read per plate as representative of 100% and background absorbance was subtracted. Three independent replicates of the assay were repeated per cell line.

### Statistical Analysis

Statistical analysis was performed in GraphPad Prism (RRID:SCR_002798), version 9.3.1 (GraphPad Software, Inc, La Jolla, CA). A p-value of <0.05 was considered statistically significant. The area under tumor growth curves was compared by one-way ANOVA and Holm-Sidak post hoc test was used to correct for multiple comparison of type I error.

## Results

### Chimeric VMG exhibits cytopathic effects across multiple sarcoma subtype cell lines

We synthesized our chimeric virus (VMG) by insertion of the MorV G gene and intergenic regions into the backbone of pVSV-XN2, replacing the VSV glycoprotein (G) gene (Fig.1A). The resultant engineered virus VMG had full replication ability and was used to infect human, murine, and canine cell lines representing a diversity of sarcoma subtypes (Table 1). We found that all tumor cell lines showed susceptibility to oncolysis especially at multiplicity of infection (MOI) 10 and MOI 1 after 72 hours (h) in culture (Fig. 1B). All cell lines showed a percentage of cell death which was dose dependent on the initial MOI administered. Only one human cell line (SK-LMS-1) exhibited resistance to cell death, showing greater than 50% viability after 72 hours post infection (hpi) at an MOI of 0.1 VMG. With an *in vitro* infection MOI of 10, the canine sarcoma line OSCA-71 exhibited the highest viability 72 hpi with a mean viability of 15% by MTS assay. The remaining live adherent cells after infection with a low MOI (0.1) were also visualized by crystal violet staining (Fig.1C). Viability assessment by MTS and crystal violet methodologies agreed for 6 out of the 7 adherent cell lines tested. In the crystal violet assay, SK-LMS-1 and 143B were the only lines with live adherent cells remaining after infection, while OSCA-71 was shown to have complete loss of adherent cells. We also engineered a GFP-expressing VMG with which all sarcoma cell lines showed infection and varying degrees of GFP production after 24 h (Supp.Fig.1).

**Table 1:**
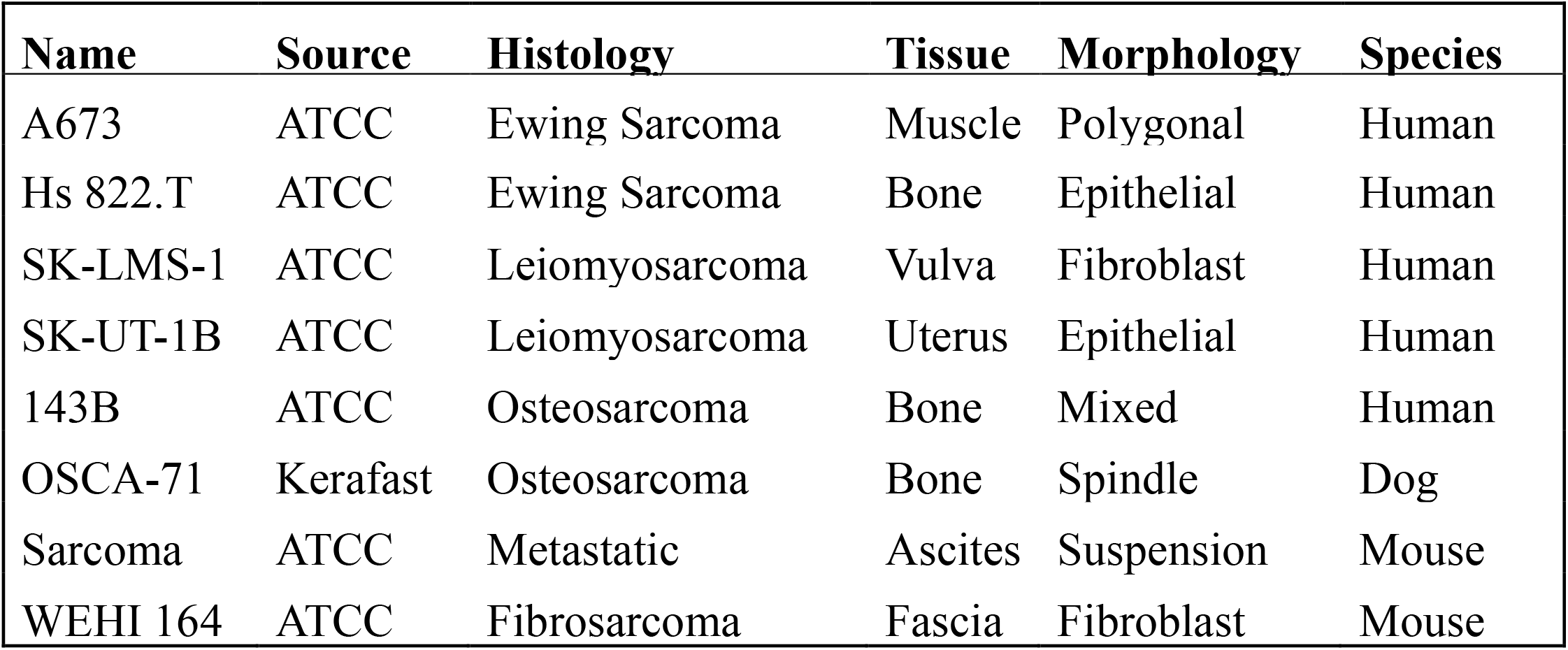
Sarcoma cell lines used, including their histology, tissue, and species of origin

| **Name** | **Source** | **Histology** | **Tissue** | **Morphology** | **Species** |
| --- | --- | --- | --- | --- | --- |
| A673 | ATCC | Ewing Sarcoma | Muscle | Polygonal | Human |
| Hs 822.T | ATCC | Ewing Sarcoma | Bone | Epithelial | Human |
| SK-LMS-1 | ATCC | Leiomyosarcoma | Vulva | Fibroblast | Human |
| SK-UT-1B | ATCC | Leiomyosarcoma | Uterus | Epithelial | Human |
| 143B | ATCC | Osteosarcoma | Bone | Mixed | Human |
| OSCA-71 | Kerafast | Osteosarcoma | Bone | Spindle | Dog |
| Sarcoma | ATCC | Metastatic | Ascites | Suspension | Mouse |
| WEHI 164 | ATCC | Fibrosarcoma | Fascia | Fibroblast | Mouse |

**Figure 1:**
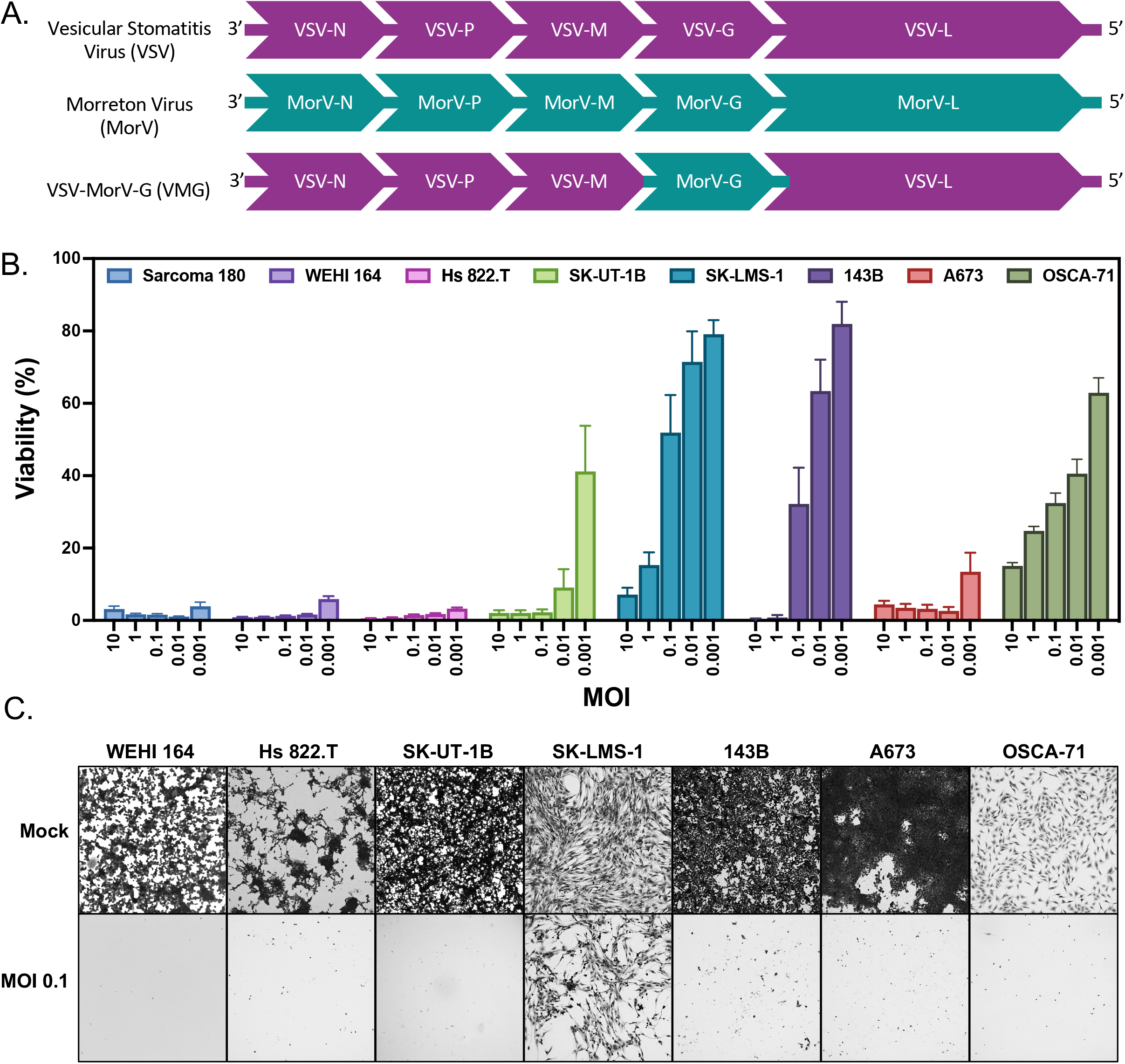
VMG genome and cytotoxicity evaluation. A) Hybrid virus construct. Morreton G protein and intergenic regions were inserted into VSV genome. B) Cytotoxicity measured by MTS (Promega). Cells were seeded at 1e4/well in 96-well plates and rested overnight. Viral dilutions were added at the indicated MOI. After 72 h, MTS was added to each well and absorbance was measured on a Cytation3 plate reader. Reads were normalized by highest absorbance per cell line as representative of maximum cell viability with background subtracted. Data was collected from three replicates per experiment over at least three independent experiments. Bars indicate mean ± SEM of n=9 data points. C) Crystal violet staining. Cells were plated at 2e5/well in 12-well plate and rested overnight. The following day they were infected with VMG at an MOI of 0.1. Cells were fixed and stained with crystal violet 72 h post-infection and phase images were captured at 10x magnification on an EVOS FL Auto Imaging System. Images are representative of experiments performed in duplicate.

### Heterogeneity of responses to VMG-induced cell death are not explained by receptor expression, progeny production, or IFNβ production

We next sought to explain differences in sarcoma cell viability after VMG infection and possible reasons for incomplete oncolysis with a replicating virus. Although the receptor for MorV has not been defined, we explored the possibility of the primary cellular receptor of VSV, LDLR, expression determining entry of virus across cell lines tested. Cell surface expression of LDLR did not correlate with proportion of cell death (Supp.Fig.2). Defects in viral entry were further ruled out by performing viral kinetics assays where an MOI of 0.1 resulted in infectious titers of comparable orders of magnitude (10^7^-10^9^ TCID_50_) by 24 hpi, as quantified in cell supernatant (Supp.Fig.3). Cell lines A673 and Hs822.T produced IFNβ *in vitro* at 48 hpi with VMG, but only at MOI 1 and not with MOI 10. Importantly, IFNβ production did not correlate with the varied percentages of oncolysis observed after infection with VMG (Supp.Fig.4).

### VMG exhibits an intermediate rate of apoptosis induction between VSV and MorV at equivalent MOI

To investigate real-time rates of infection and killing by VSV, MorV, and VMG, we utilized the IncuCyte S3 system with an Annexin V Red fluorescent reporter for apoptotic cell death. Phase and red fluorescence images of the A673 cell line were taken every 3 h for a total of 72 h after introducing virus and the red area was used to quantify the relative level of apoptosis per viewing field at each timepoint. Values were plotted for each virus and MOI to generate non-linear regressions representative of viral killing (Fig.2A,C). The highest MOI of each virus reached maximum rate of apoptosis at the earliest timepoint. From the best-fit curves, the time at which 50% of the maximal cell death occurred could be determined (T_50_) as a quantitative measure of efficacy between MOIs and across viruses (Fig. 2B,D). This data correlates with the viral kinetics (Supp.Fig.3), in which progeny production at MOI 0.1 peaks at 24 h for A673, immediately prior to a dramatic increase in apoptotic cell death as observed in the IncuCyte Annexin V assay. Additionally, VMG exhibited an intermediate T_50_ value between VSV and MorV in the A673 cell line.

**Figure 2:**
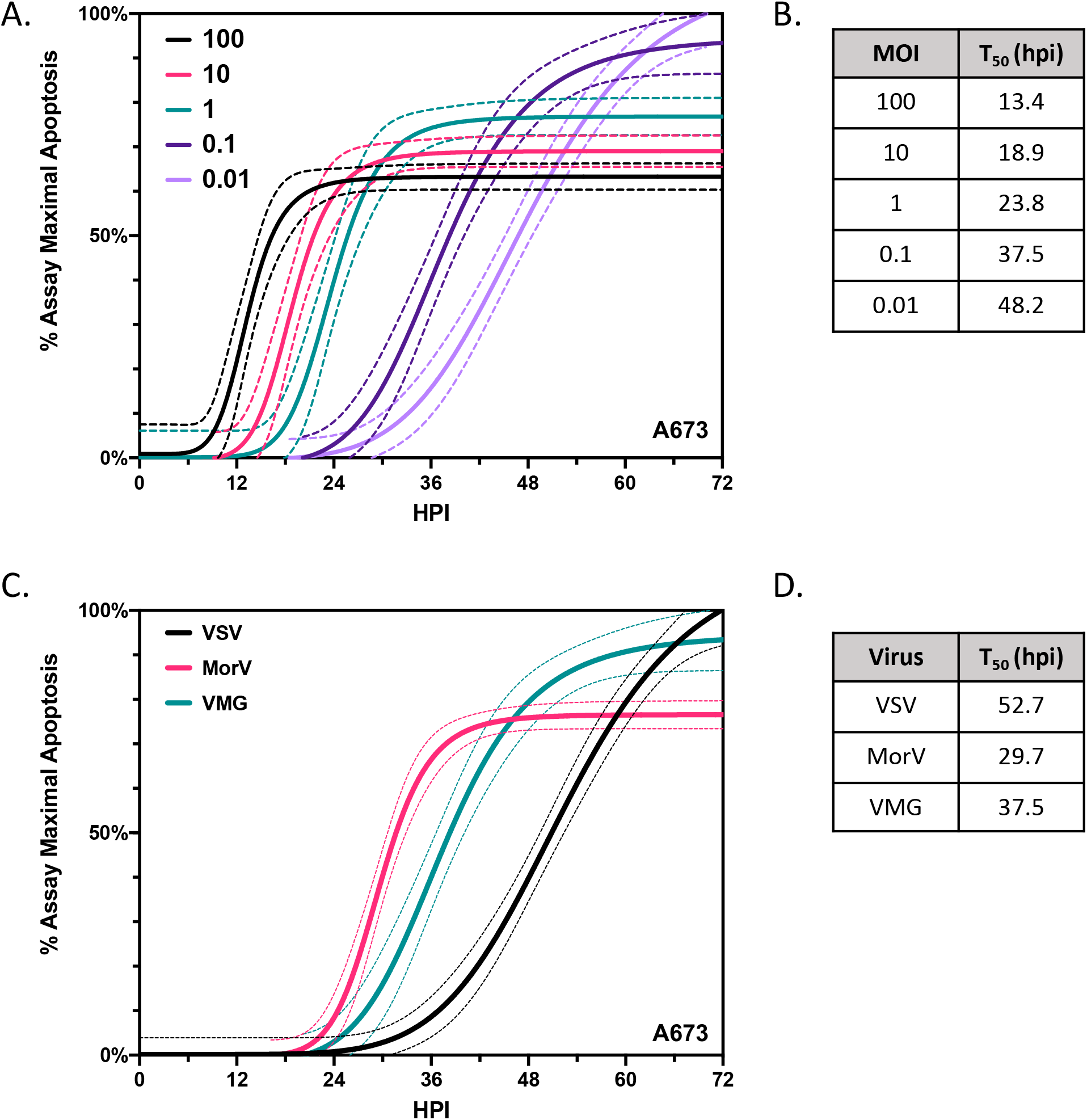
Apoptosis quantification in real-time. A) VMG induced apoptosis *in vitro* in A673 Ewing sarcoma cell line. 1e4 A673 cells per well were plated in a 96-well plate and rested overnight. The next day, Annexin V Red (Essen BioScience) solution was added to experimental wells and viral dilutions were added in three replicate wells per MOI. Four images per well were taken every 3 hours in the IncuCyte S3 system (Essen BioScience) for 72 h, collecting both phase and red fluorescence data. Red object area for each image was quantified, background subtracted for each timepoint, and normalized to the maximum red area for the entire experiment. Data were graphed and standard curves were generated using a sigmoidal, 4PL interpolation with a 95% confidence interval. Best fit values gave a T_50_ value, the time at which 50% of cells have undergone apoptosis. C) Comparison of VSV, MorV, and VSV-MorV-G cytotoxicity in A673 cells at MOI 0.1 analyzed in the IncuCyte S3 System. D) Interpolated timepoint at which 50% of maximal cell death occurs compared between viruses from the data in C.

### Intranasal administration of VMG is well tolerated in an immunocompetent mouse model

To assess whether infection with VMG is associated with neurotoxicity in laboratory mice as shown for VSV, we intranasally administered VMG at four different doses (10^7^, 10^8^, 10^9^, or 10^10^ TCID_50_ in 10 microliters total volume) to immunocompetent mice. Short term (3 days) and long term (~7 weeks) toxicity was evaluated as previously described.^9^ Overall, all doses were well tolerated. In the long-term observation group, only one mouse experienced a drop in body weight (<20% reduction) at the highest dosage of VMG before recovering (Supp.Fig.5).

### Complete tumor inhibition is observed after a single administration of VMG in a xenograft sarcoma model

To determine if VMG can delay tumor growth in a xenograft model of Ewing sarcoma (A673), we treated tumor-bearing female NOD-SCID mice intratumorally (IT) at two doses (2e^5^ or 2e^6^ TCID_50_) of VMG. In comparison to the vehicle group, there was significant tumor inhibition (p<0.0001) after IT administration with a single dose of VMG at either of the two dosages (Fig.3A). However, adverse events were observed in the VMG treated groups, with one mouse found moribund on day 6, and 6 mice found dead after day 8 (Fig.3B). Liver function tests were performed on serum samples collected from terminal bleeds for albumin (ALB), aspartate transaminase (AST), alanine aminotransferase (ALT), alkaline phosphatase (ALP), total bilirubin, and glucose (GLU) (Fig.3C). We found that the one moribund mouse had the highest level of ALT and serum levels of ALB and ALP increased in a dose-dependent manner 15 days after treatment with VMG, indicating possible liver toxicity in the immunodeficient model.

**Figure 3:**
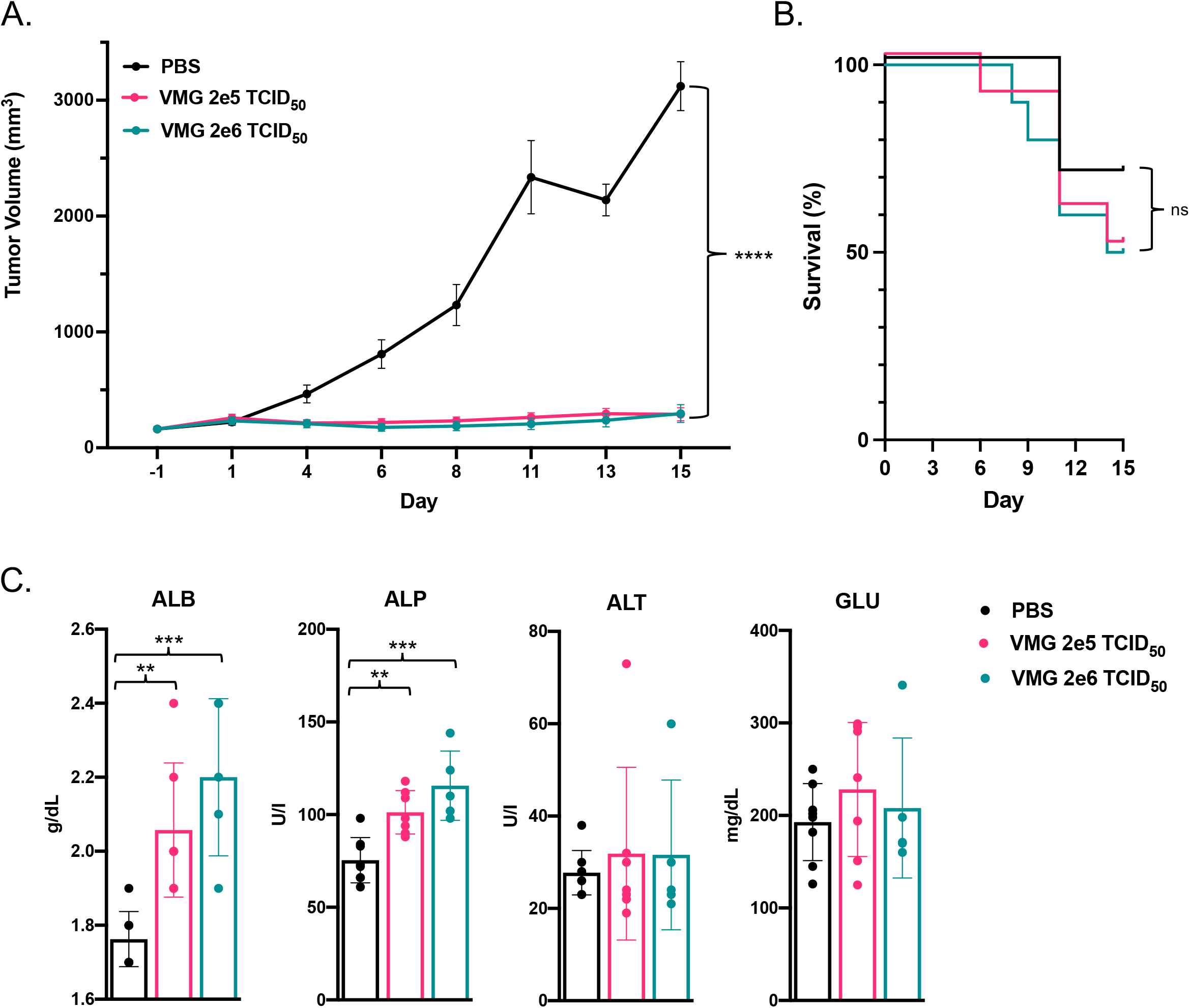
Efficacy of VMG in xenograft sarcoma model A673. A) Average tumor volume of 10 female mice per group. When average tumor volume was over 150 mm3, mice received intratumoral injection with either PBS, VMG at a dose of 2e5 TCID50, or VMG 2e6 TCID50 (Day 0). Tumor volume was measured three times weekly until a humane endpoint, death, or end of study (Day 15). Data plotted as mean +/−SD. Area under tumor growth curves was compared by one-way ANOVA with Holm-Sidak correction for type I error. B) Overall survival per group. Endpoints included humane sacrifice due to tumor size or ulceration, or death. C) Serum levels of liver enzymes. Liver function tests were performed with serum taken at study endpoint. Significance was calculated using one-way ANOVA. **p<0.005 ***p<0.0005 ****p<0.00005.

### Intratumoral injection of VMG results in increased CD8 T cell infiltration in a syngeneic model of fibrosarcoma

We next established an immunocompetent murine model of fibrosarcoma. WEHI 164 tumors were established subcutaneously in male and female Balb/c mice. Three doses of 10^7^ TCID_50_ were administered IT at days 0, 5, and 10. No significant difference in tumor volume between the PBS control and VMG treated groups was observed by end of study (Fig.4A). At day 14, mice were sacrificed, and tumor tissue collected if visible. Single-cell suspensions were isolated from freshly collected murine tumor tissue and viably cryopreserved. Tumor infiltrating lymphocyte (TIL) subsets were evaluated by with an 18-color flow cytometry panel (Table 2). Live CD45 cells were analyzed for the frequencies of immune cell types and compared between the control group and VMG treated mice. Notably, the proportion of CD3+ cells within the CD45 compartment increased after VMG treatment (Fig.4B). Within the CD3 compartment, CD8+ cytotoxic T cells were significantly increased (Fig.4C). The CD45 compartment within each tumor was composed of primarily CD11b+ cells, although the overall proportion of CD11b+ cells decreased after treatment with VMG (Fig.4D). Additionally, the percentage of Tregs (FoxP3+CD25+) within the CD4 compartment decreased within several of the VMG treated mice but did not reach statistical significance (Fig.4E). We performed unbiased clustering analysis by t-distributed stochastic neighbor embedding (t-SNE), which confirmed an increase of the CD3+ compartment in VMG-treated tumors, especially CD8+ cells, and an overall decrease in the CD11b+ compartment (Fig. 4F,G).

**Table 2:** Fluorescent antibody panel for WEHI 164 immunophenotyping

| **mAb** | **Fluor** | **Clone** | **Supplier** | **Catalog #** |
| --- | --- | --- | --- | --- |
| Ly6G | BB700 | 1A8 | BD | 566453 |
| F4/80 | PerCP | BM8 | BioLegend | 123125 |
| CD19 | AF488 | 6D5 | BioLegend | 115524 |
| CD11c | BV786 | HL3 | BD | 563735 |
| PD-1 | BV711 | 29F.1A12 | BioLegend | 135231 |
| CD45 | BV650 | 30-F11 | BioLegend | 103151 |
| CD3 | BV605 | 17A2 | BioLegend | 100237 |
| CD335 (NKp46) | BV510 | 29A1.4 | BioLegend | 137623 |
| FoxP3 | BV421 | MF23 | BD | 562996 |
| CD4 | BUV805 | GK1.5 | BD | 612900 |
| Viability | Zombie UV |  | BioLegend | 423107 |
| CD8 | BUV395 | 53-6.7 | BD | 565968 |
| I-A/I-E (MHC-II) | APC-Cy7 | M5/114.15.2 | BioLegend | 107627 |
| Ki67 | AF700 | 16A8 | BioLegend | 652419 |
| CD25 | APC | PC61 | BioLegend | 102011 |
| CD11b | PE-Cy7 | M1/70 | BioLegend | 101215 |
| Ly6C | PDzl594 | HK1.4 | BioLegend | 128043 |
| CD49b | PE | DX5 | BioLegend | 108907 |

**Figure 4:**
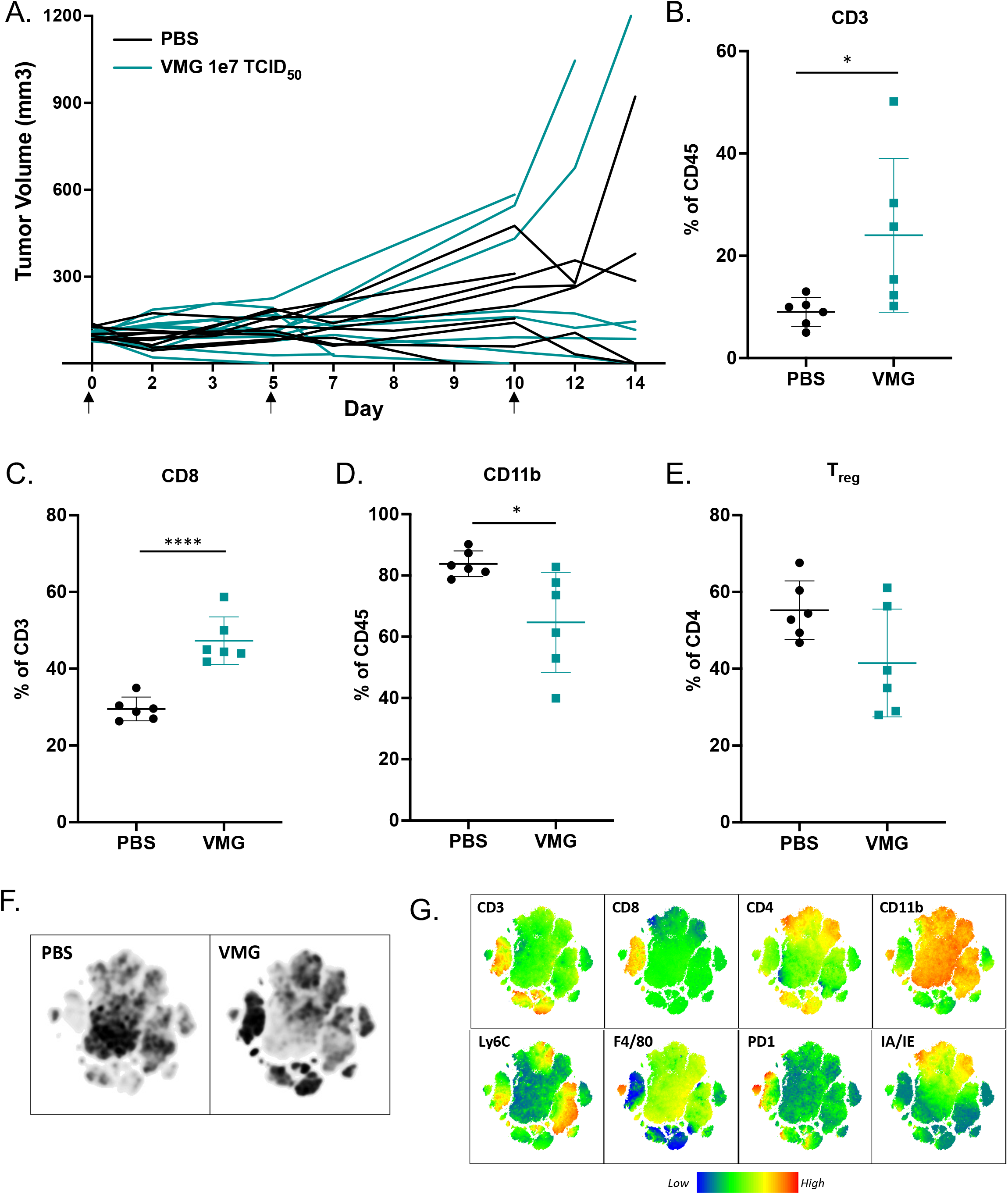
Assessment of VMG efficacy in syngeneic model of sarcoma WEHI 164. A) Individual murine tumor volumes after treatment. BALB/c mice were inoculated subcutaneously with WEHI 164. When tumor volume was between 80-120 mm3, mice were randomly divided into 2 groups and received intratumoral injection with either PBS (n=9) or VMG (n=11) at a dose of 1e7 TCID_50_ (Day 0). Injections were repeated at day 5 and 10. Tumor volume was measured three times weekly until humane endpoint or end of study (Day 14). B-E) Tumor infiltrating lymphocytes (TILs) were compared between VMG and PBS control mice by flow cytometry. Results were analyzed by unpaired t-test. *p<0.05, ****p<0.0001 F) t-distributed stochastic neighbor embedding (t-SNE) analysis performed on live CD45+ TILs. G) Heatmap statistic for t-SNE plots, indicating areas of low4/no expression (blue) and highest expression (red) for the indicated cell markers

### Mouse IFNβ inhibits VMG cell killing in WEHI 164 in a dose-dependent manner

To investigate possible explanation for a lack of tumor inhibition in the WEHI 164, we tested whether exogenous murine IFNβ (mIFNβ) could protect WEHI 164 from VMG-induced cell death. Non-cancer cells within the tumor microenvironment could mount an anti-viral response after VMG administration, leading to inhibition of viral killing if the WEHI 164 cancer cells are responsive to IFNβ. Our results showed that WEHI 164 cells were highly responsive to IFNβ, and at an MOI of 1, a concentration of 500 U/mL IFNβ rescued approximately 50% of cells from VMG-induced cell death (Fig.5A). There was also dose-dependent cell death induced by highest concentrations of mIFNβ alone. The cell line utilized for the xenograft model also was protected from virally induced cell death by exogenous human IFNβ (Fig.5B). From the 8 cell lines tested, 4 maintained responsiveness to IFNβ which abrogated cell death post-infection, while 4 cell lines exhibited no protective responses with type I IFN co-culture (Supp.Fig.6).

**Figure 5:**
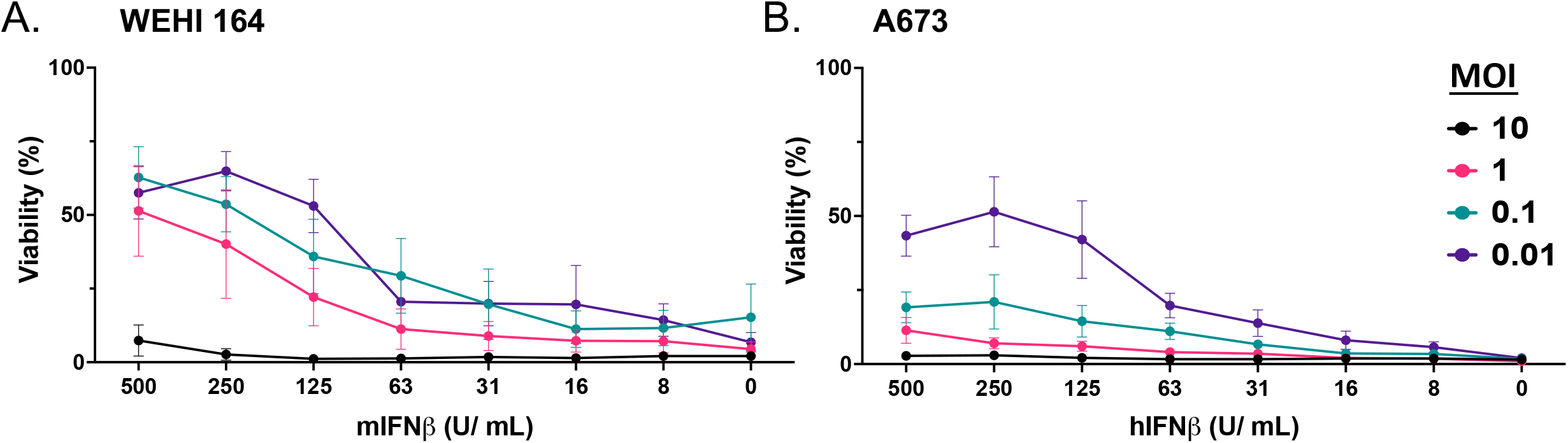
Effect of exogenous IFNβ on viral killing. Dose and MOI dependent responses to species-specific exogenous species-specific IFNβ on sarcoma cell lines used as tumor models A) WEHI-164 and B) A673. Data points are three independent experimental replicates plotted as mean ± SEM.

## Discussion

In this study, we show that the chimeric virus, VMG, exhibited oncolysis in all sarcoma cell lines at high MOI (10 and 1), with variable cytotoxic effect at lower MOI (0.1 and 0.01) *in vitro*. The metabolic cell viability assay yielded higher viability than indicated visually with crystal violet staining, demonstrating the importance of multiple assessment methods to determine cellular viability.

Notably, detectable GFP expression 24 hours after infection with VMG carrying the GFP transgene did not correlate with viability outcomes at 72 hours, indicating that cells present at early infection timepoints may have phenotypic differences affecting their sensitivity to infection or virally induced cell death. Heterogeneity within cancer cell lines prior to infection may have affected susceptibility to infection or killing. Differences observed in cell viability at 72 hpi were not due to LDLR expression, and the definitive receptor or receptor family for MorV remains to be determined. The MorV G protein could use another LDLR family member or another cell surface entry target, given the genetic difference between sequences of MorV G and VSV G. However, receptor availability is unlikely to be the cause of variable cell death between cell lines since the virus infected and replicated across all sarcoma cell lines tested, as evidenced by similar titers of infectious VMG 24 hpi with MOI 0.1. *In vitro*, human sarcoma cell lines did not produce detectable levels of IFNβ in response to viral infection. This result is not unexpected, as the VSV M protein is known to inhibit type I IFN production, and remains intact in our hybrid virus.^10^ The most resistant cell line, SK-LMS-1, is a sarcoma line tested derived from vulvar tissue. Further work is needed to characterize this cell line’s inherent mode of resistance to VMG cell killing.

The A673 cell line was highly sensitive to viral killing *in vitro* and was used to generate a xenograft model to test intratumoral efficacy of VMG. Complete tumor inhibition was observed in this immunodeficient model, but some adverse events were observed, hypothesized to be mainly due to liver damage based on elevated serum liver-associated enzymes. These data should also be evaluated in the context of the model being a highly immunodeficient one. Correspondingly, we have shown that intranasal administration of VMG was well tolerated in an immunocompetent mouse model, with a drop in body weight only associated with the highest tested dose (10^10^ TCID_50_)

Despite the oncolytic success of VMG in the A673 model, there was little to no observable reduction in tumor growth in the WEHI 164 murine fibrosarcoma model. Although WEHI 164 showed significantly reduced viability in the MTS assay after VMG infection, tumor growth was not affected in the subcutaneous model after 3 doses of virus. However, we showed an appreciable difference in tumor infiltrating lymphocytes between the PBS control group and VMG treated tumors, namely an increase in CD8+ T cells. Future endeavors will be focused on elucidating whether this expanded population of cytotoxic T cells within the treated tumor are tumor-or VMG-specific, or both. One possible explanation for these differing results is that WEHI 164 tumor cells could have responded to type I IFN produced by stromal cells in the microenvironment and entered an anti-viral state. This could have slowed viral replication down to tip the balance in favor of cancer cell replication. In contrast, the human cell line A673 would not have been protected by mouse IFNβ *in vivo* due to species interferon receptor (IFNAR) specificity. Another possible factor for the lack of effect in WEHI 164 could have been immune clearance of the virus before substantial oncolysis had taken place. The mechanism of this *in vivo* resistance remains to be clarified, but the first hypothesis that IFN could have limited viral replication and cell death is supported by the follow-up *in vitro* experiments testing the effect of exogenous IFNβ during infection on WEHI 164 cells. Although we did not observe a tangible difference in tumor volume within the WEHI 164 model, the altered immune landscape after oncolytic virotherapy may still offer therapeutic benefit when combined with other immunotherapies. Clinical relevance is shown in human Ewing sarcoma, where a high frequency of T cells and activated natural killer (NK) cells correlated with prolonged overall survival.^11^ These results show VMG does elicit an immune response within a murine sarcoma model. Our data also shows that cancer cell lines vary in their ability to produce or respond to exogenous IFNβ. Therefore, it is important to characterize patient-specific level of IFN pathway defects within a tumor prior to administering vesiculovirus-based oncolytics. Future work will aim to elucidate mechanisms of resistance to oncolytic virotherapy, including the possibility of protection by local type-I IFN produced by non-cancerous cells.

Overall, we show that VMG exhibits oncolysis in multiple sarcoma subtypes and can stimulate CD8+ T cell infiltration in a syngeneic model of fibrosarcoma. This hybrid virus can advance viral therapy as a potential safer alternative to VSV without viral attenuation through an IFNβ transgene. We conclude that VMG should be evaluated as a novel oncolytic platform in sarcoma. Future preclinical studies will focus on improved viral engineering, toxicity studies, and assessment of combination immunotherapies.

## Abbreviations

ALB: albumin
ALP: alkaline phosphatase
ALT: alanine aminotransferase
AST: aspartate aminotransferase
CPE: cytopathic effect
FBS: fetal bovine serum
GFP: green fluorescent protein
GGT: gamma-glutamyl transferase
GLU: glucose
h: hours
hIFNβ: human interferon beta
hpi: hours post-infection
IFNβ: interferon beta
IFNAR: interferon alpha receptor
IGR: intergenic regions
IT: intratumoral
mIFNβ: murine interferon beta
MOI: multiplicity of infection
MorV: Morreton virus
MTD: maximum tolerated dose
MTS: [3-(4,5-dimethylthiazol-2-yl)-5-(3-carboxymethoxyphenyl)-2-(4-sulfophenyl)-2H-tetrazolium, inner salt
NOD: SCIDnon-obese diabetic severe combined immunodeficient NK cells natural killer cells
NU/J: athymic nude mouse OS overall survival
OV: oncolytic virus
RPMI: Roswell Park Memorial Institute medium
SD: standard deviation
TBIL: total bilirubin
TCID50: 50% tissue culture infectious dose
t-SNE: t-distributed stochastic neighbor embedding
VMG: VSV-MorV-G
VSV: Vesicular stomatitis virus

## Acknowledgments

We thank the personnel of the Department of Comparative Medicine at Mayo Clinic in Scottsdale, Arizona for their assistance during animal studies. This work was supported by the National Institute of Health (NIH) through a K01 award CA234324 (to B.M.N.), Marley Endowment Funds (Mayo Clinic Arizona) (to B.M.N.), National Cancer Institute (NCI) K12 award CA090628 (to M.J.B.), Hepatobiliary SPORE grant (to M.J.B.), Mayo Clinic Cancer Center CCSG Gene and Virus Therapy program grant (to M.J.B.), P50 grant CA210964 (to M.J.B. and B.M.N.), and a Department of Molecular Medicine (Mayo Clinic) Small Grant (to C.D.).

## Author Contributions

Conceptualization, C.D., M.J.B., B.M.N.; Methodology, C.D., O.B., A.T.B., Y.Z., M.J.B., and B.M.N.; Investigation, C.D.; Visualization, C.D.; Writing – Original Draft, C.D.; Writing-Review & Editing, C.D., M.S., O.B., N.M.E., A.T.B., Y.Z., M.G., E.R., M.T.B., K.H.B., B.J., M.J.B., B.M.N; Funding Acquisition, C.D., M.S., M.J.B., B.M.N; Resources, M.J.B. and B.M.N.; Supervision, M.J.B. and B.M.N.

## References

Gage MM, Nagarajan N, Ruck JM, Canner JK, Khan S, Giuliano K, Gani F, Wolfgang C, Johnston FM, Ahuja N, Gage MM, Nagarajan N, Ruck JM, Canner JK, Khan S, Giuliano K, Gani F, Wolfgang C, Johnston FM, Ahuja N. Sarcomas in the United States: Recent trends and a call for improved staging. Oncotarget. 2019;10(25):2462–2474. doi:10.18632/ONCOTARGET.26809

Charles Hesla A, Papakonstantinou A, Tsagkozis P. cancers Current Status of Management and Outcome for Patients with Ewing Sarcoma. Cancers (Basel). 2021;13:1202. doi:10.3390/cancers13061202

Birdi HK, Jirovec A, Cortés-Kaplan S, Werier J, Nessim C, Diallo JS, Ardolino M. Immunotherapy for sarcomas: new frontiers and unveiled opportunities. J Immunother Cancer. 2021;9:1580. doi:10.1136/jitc-2020-001580

Paglino JC, van den Pol AN, Klose C, Berchtold S, Schmidt M, Beil J, et al. Biological treatment of pediatric sarcomas by combined virotherapy and NK cell therapy. Sarcoma. 2019;12(1):2279–2288. doi:10.1186/s13045-019-0756-z

Felt SA, Grdzelishvili VZ. Recent advances in vesicular stomatitis virus-based oncolytic virotherapy: A 5-year update. J Gen Virol. 2017;98(12):2895–2911. doi:10.1099/jgv.0.000980

Johnson JE, Nasar F, Coleman JW, Price RE, Javadian A, Draper K, et al. Neurovirulence properties of recombinant vesicular stomatitis virus vectors in non-human primates. Virology. 2007;360(1):36–49. doi:10.1016/j.virol.2006.10.026

Kelley JM, Emerson SU, Wagner RR. The Glycoprotein of Vesicular Stomatitis Virus Is the Antigen That Gives Rise to and Reacts with Neutralizing Antibody. J Virol. 1972;10(6):1231. doi:10.1128/jvi.10.6.1231-1235.1972

Brun J, McManus D, Lefebvre C, Hu K, Falls T, Atkins H, et al. Identification of genetically modified maraba virus as an oncolytic rhabdovirus. Mol Ther. 2010;18(8):1440–1449. doi:10.1038/mt.2010.103

Nagalo, BM, Zhou, Y, Loeuillard, EJ, Dumbauld, C, Barro, O, Elliott, NM, et al. Characterization of Morreton Virus (MORV) as a Novel Oncolytic Virotherapy Platform for Liver Cancers. Hepatology. 2022. doi:10.1002/HEP.32769

Faul EJ, Lyles DS, Schnell MJ. Interferon response and viral evasion by members of the family rhabdoviridae. Viruses. 2009;1(3):832–851. doi:10.3390/v1030832

Stahl D, Gentles AJ, Thiele R, Gütgemann I. Prognostic profiling of the immune cell microenvironment in Ewings Sarcoma Family of Tumors. Oncoimmunology. 2019;8:12. doi:10.1080/2162402X.2019.1674113

Lawson ND, Stillman EA, Whitt MA, Rose JK. Recombinant vesicular stomatitis viruses from DNA. Proc Natl Acad Sci U S A. 1995;92(10):4477–4481. doi:10.1073/PNAS.92.10.4477

