## Supplemental Figures for "Multi-modal Efficacy of a Chimeric Vesiculovirus Expressing the Morreton Glycoprotein in Sarcoma"

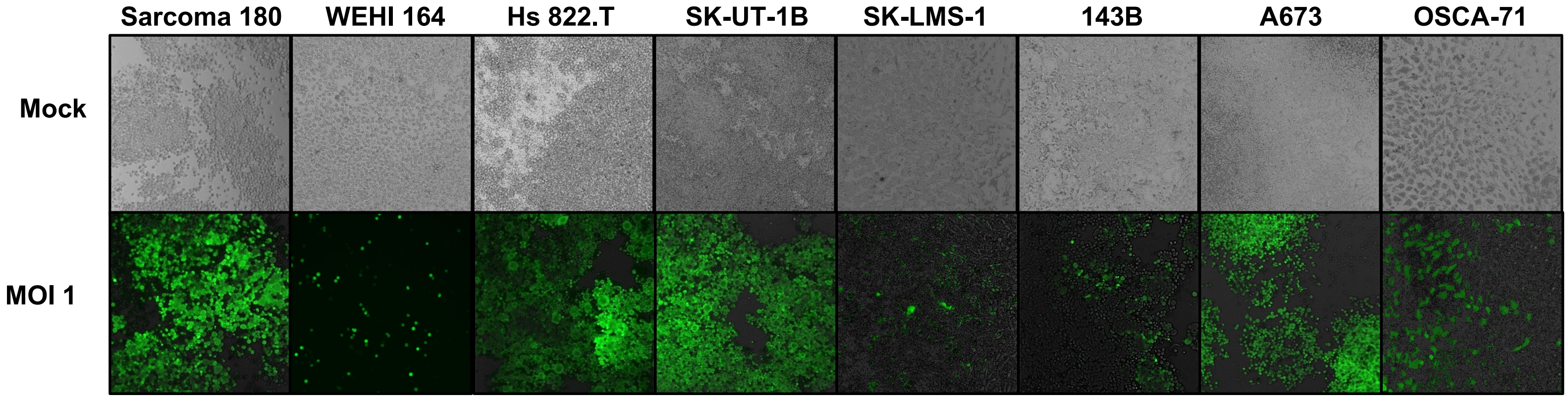


Supplemental Figure 1: *VMG-GFP production 24 h after infection with MOI 1.* Each cell line was infected for 24 h in a 6-well plate with VMG carrying the GFP transgene. Phase and green fluorescent images were taken at 10x magnification using an EVOS FL Auto microscope. Representative images are taken from experiments completed in duplicate.

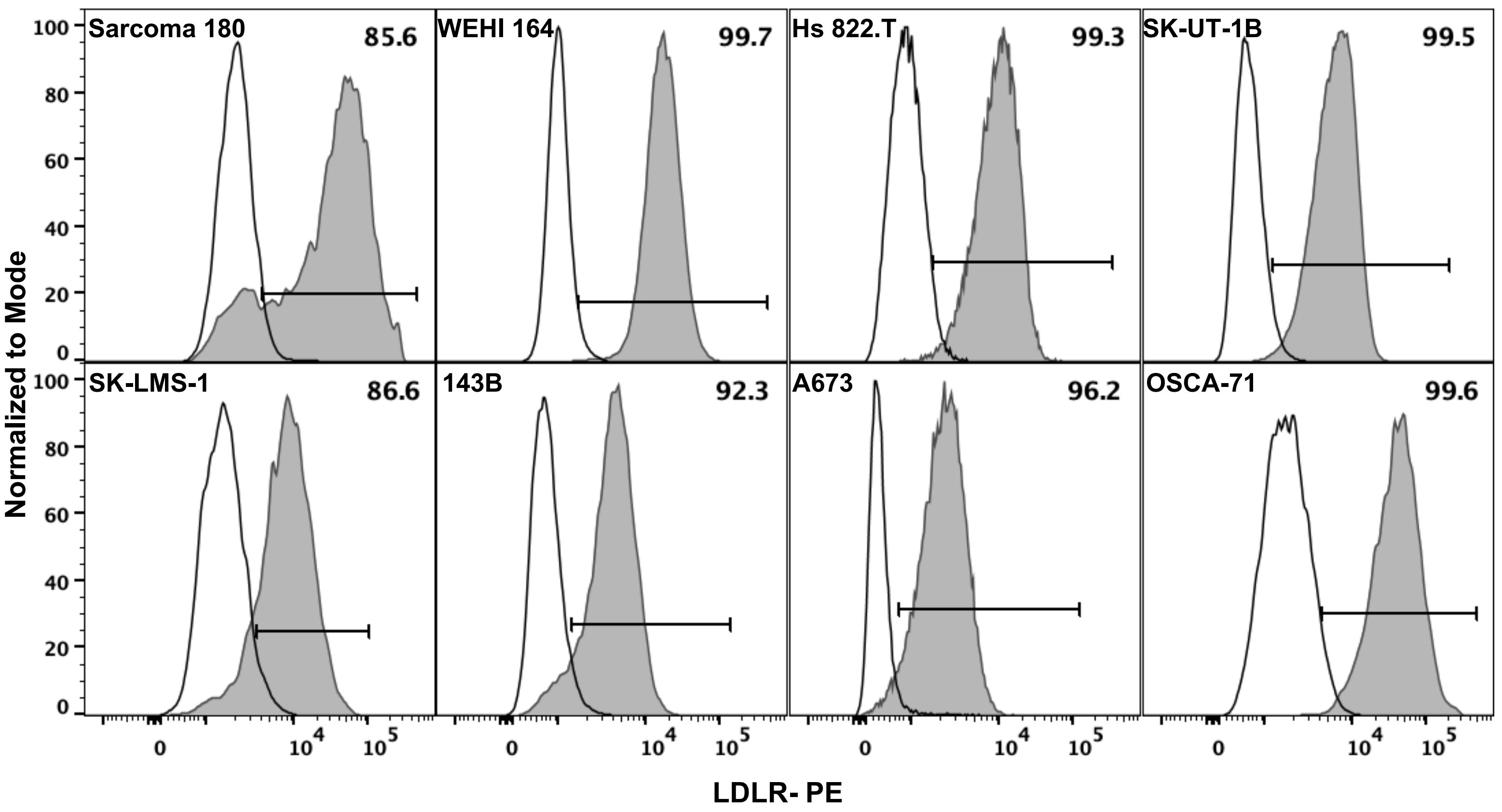


Supplemental Figure 2: Species-specific* LDLR Expression of sarcoma cell lines (shaded histogram) gated on 5% isotype control (unshaded histogram). OSCA-71 was stained with anti-mouse LDLR due to cross-reactivity of the antibody. Representative histograms are from experiments repeated in duplicate.

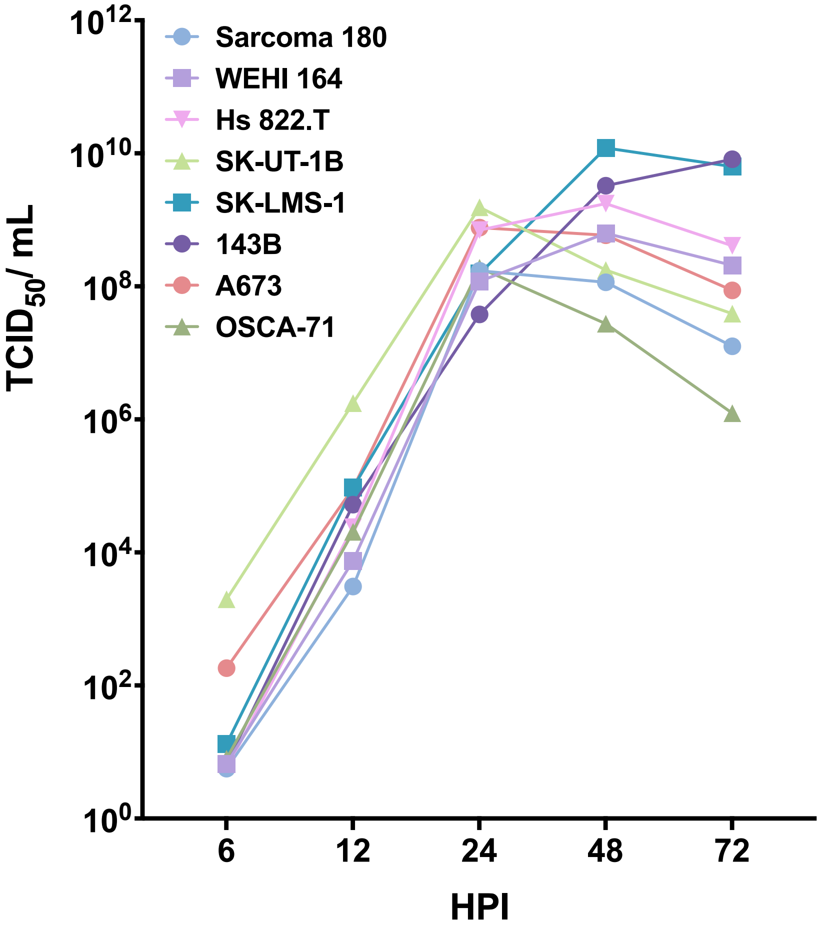


Supplemental Figure 3: *Viral growth kinetics in sarcoma cell lines.* Cells were plated in 6-well plates at 2.5e5/well and rested overnight. The next day they were infected with VMG at MOI of 0.1. One hour post-infection, media was replaced. At each timepoint, supernatant was collected. Fifty percent tissue culture infectious dose ($TCID_{50}$) values were determined by the Spearman-Kärber algorithm using serial dilutions in BHK-21 cells. Data are plotted from at least three independent experiments quantifying $TCID_{50}$ for each point with mean ± SEM.

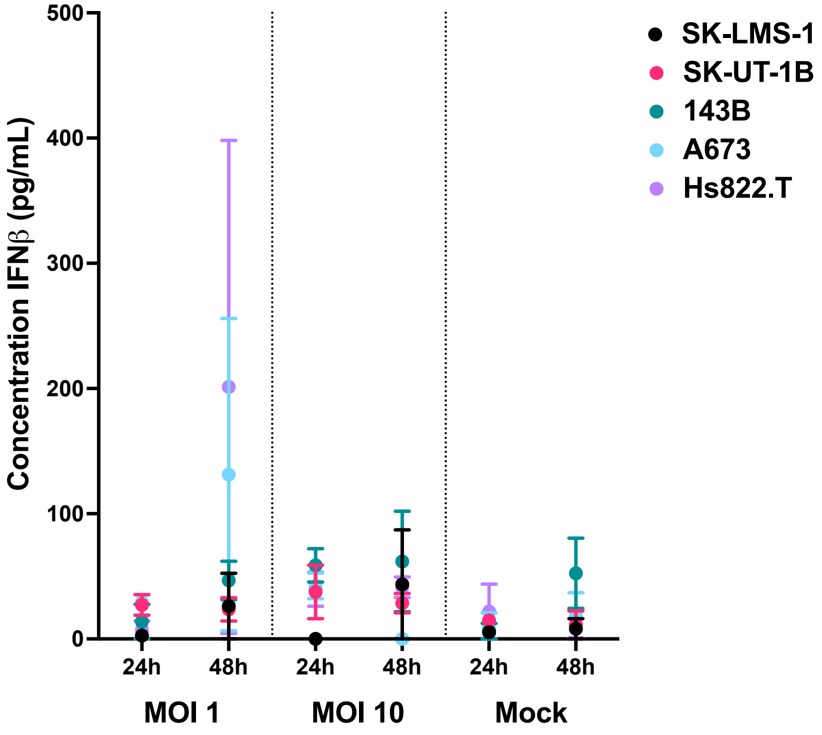


Supplemental Figure 4: *IFN$\beta$ production of human sarcoma cell lines.* Cells were treated with VMG at MOI 10 and MOI 1 and supernatant collected at 24 and 48 HPI as assessed by ELISA. Data is plotted as mean $\pm$ SD per condition, with two technical replicates per condition.

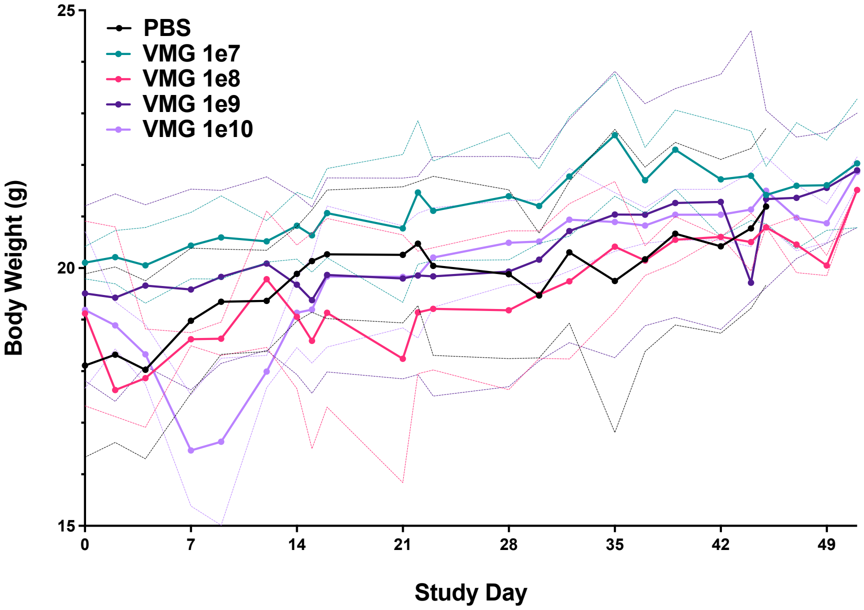


Supplemental Figure 5: *Monitoring of body weight following intranasal administration VMG*. Body weight was recorded every 2-3 days until end of study. n=7 mice per group. Mean ± SD per condition are plotted.

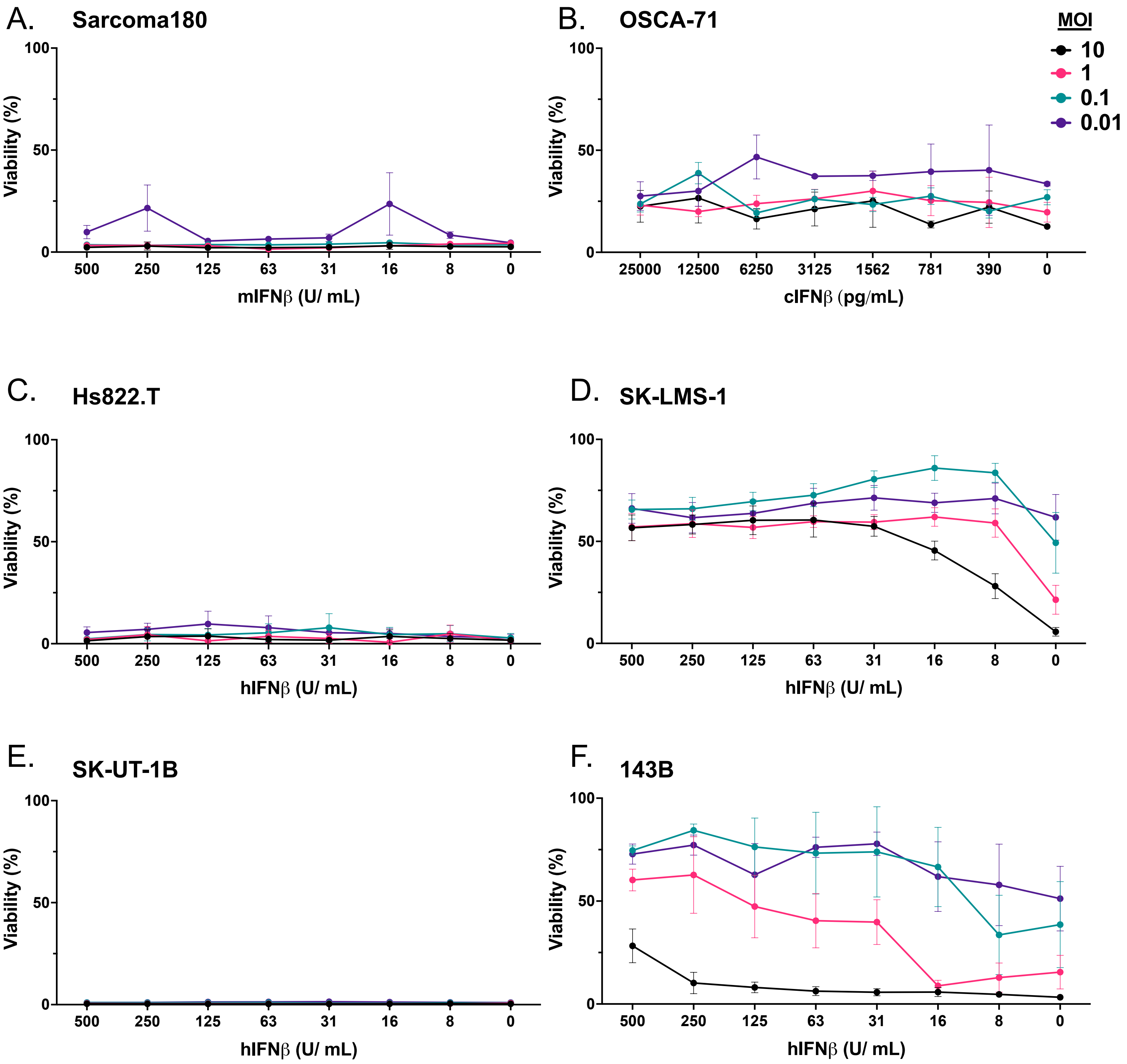


Supplemental Figure 6: *Dose- and MOI-dependent responses to species-specific exogenous IFNβ in sarcoma cell lines*. Data points are representative of mean ± SEM from three independent experimental replicates.
